# A natural variant of the essential host gene *MMS21* restricts the parasitic 2-micron plasmid in *Saccharomyces cerevisiae*

**DOI:** 10.1101/2020.06.11.146555

**Authors:** Michelle Hays, Janet M. Young, Paula Levan, Harmit S. Malik

## Abstract

Ongoing antagonistic coevolution with selfish genetic elements (SGEs) can drive the evolution of host genomes. Here, we investigated whether natural variation allows some *Saccharomyces cerevisiae* strains to suppress 2-micron (2μ) plasmids, multicopy nuclear parasites that have co-evolved with budding yeasts. To quantitatively measure plasmid stability, we developed a new method, Single-Cell Assay for Measuring Plasmid Retention (SCAMPR) that measures copy number heterogeneity and 2μ plasmid loss in live cells. Next, in a survey of 52 natural *S. cerevisiae* isolates we identified three strains that lack endogenous 2μ plasmids and reproducibly inhibit mitotic plasmid stability. Thus, their lack of endogenous 2μ plasmids is genetically determined, rather than the result of stochastic loss. Focusing on one isolate (Y9 ragi strain), we determined that plasmid restriction is heritable and dominant. Using bulk segregant analysis, we identified a high-confidence Quantitative Trait Locus (QTL) for mitotic plasmid instability on Y9 chromosome V. We show that a single amino acid change in *MMS21* is associated with increased 2-micron instability. *MMS21* is an essential gene, encoding a SUMO E3 ligase and a member of the Smc5/6 complex, which is involved in sister chromatid cohesion, chromosome segregation, and DNA repair. Our analyses leverage natural variation to uncover a novel means by which budding yeasts can overcome a highly successful genetic parasite.

## Introduction

Host genomes are engaged in longstanding conflicts with myriad selfish genetic elements (SGEs, or genetic parasites)^1,2,3^. SGEs propagate within an organism or population at the expense of host fitness^1^. Many SGEs, including viruses, selfish plasmids, and other pathogens, must coopt the host’s cellular machinery for their own survival: to replicate their genomes, to transcribe and translate their proteins, and to ensure their proliferation by passage into new cells^1,3^. If a host variant arises that can suppress SGEs (host restriction), this variant will be favored by natural selection and can rise in frequency in a population. If resistance has no fitness cost, such variants will rapidly fix within host species. Even if these variants are slightly deleterious, such variants could be maintained in quasi-equilibrium in host species^4–7^.

Studies in diverse biological taxa have leveraged genetic mapping strategies to identify quantitative trait loci (QTL) for host resistance to parasites^8–11^. Such studies have revealed that host populations are more likely to harbor variation in resistance to coevolved, rather than to recently introduced, parasites^12^. Studying parasites in their native host context therefore maximizes opportunities to discover host resistance mechanisms. However, it is often difficult to study natural variation in resistance, because hosts and/or parasites are often intractable in the laboratory. Budding yeast provides an ideal system to study host-SGE genetic conflicts, with abundant genetic tools, together with resources for comparative and population genetics. Yeast species harbor a variety of SGEs including retrotransposable elements, RNA viruses and 2-micron plasmids^13–18^. Yet, despite its long history as a popular model eukaryote, natural variation in cellular immunity factors against SGEs has been largely uncharacterized in *S. cerevisiae* and related species^19–23^. Here, we investigated whether *S. cerevisiae* strains harbor genetic variants that allow them to resist a highly successful SGE: 2-micron plasmids.

2-micron (2μ) plasmids are nuclear SGEs found in multiple, divergent budding yeast species^24–27^. They are best characterized in *S. cerevisiae*, where they are found in high copy numbers: ~50 copies per haploid and ~100 copies per diploid cell^28,29^. Their widespread prevalence in *S. cerevisiae* and other budding yeast species has raised the question of whether 2μ plasmids might be more commensal than parasitic. In the mid-1980s, two seminal studies showed that *S. cerevisiae* strains carrying 2μ plasmids (*cir^+^)* grew 1-3% more slowly than did their *cir^0^* counterparts under laboratory conditions; thus 2μ plasmids confer a clear fitness defect^30,31^. Recent studies have reinforced the fitness defect associated with carriage of 2μ plasmids^32,33^. Furthermore, many mutant yeast strains, which are sick in the presence of the 2μ plasmids, can be partially rescued when ‘cured’ of their 2μ plasmids^34,35^. For example, *nib1* mutants (a hypomorphic allele of *ULP1*^34^) show “*nibbled*” colonies in the presence of 2μ plasmids due to colony sectoring from cells that stop dividing when overburdened with 2μ plasmids, but form smooth (wild-type) colonies in their absence^34^. These and other data^35^ suggest that 2μ plasmids impose a selective burden on yeast, both under rapid laboratory growth conditions as well as in times of stress. In contrast to bacterial plasmids, which can harbor host-beneficial ‘cargo’ genes, such as antibiotic resistance genes, no such beneficial genes have ever been observed in natural 2μ plasmids^36^. Indeed, there are no known conditions in which 2μ plasmids are beneficial to the host, further supporting that its presence is likely the result of efficient parasitism. Although they are stable in *S. cerevisiae*, experimental studies show that 2μ plasmids exhibit lower copy number and decreased stability when introduced into exogenous species^37^. These findings suggest that 2μ plasmids have co-evolved with host genomes to become a successful genetic parasite of yeasts.

To be successful, 2μ plasmids must replicate and segregate with high fidelity into daughter cells during both yeast mitosis and meiosis. Without these capabilities, plasmids risk being lost from the population as their host cells are outcompeted by plasmid-less daughter cells. Yet, 2μ plasmids encode just four protein-coding genes (represented by arrows in Figure 1A) in *S. cerevisiae*. *REP1* and *REP2* encode plasmid-encoded DNA-binding proteins that bind to the 2μ *STB* locus to mediate segregation^38–40^. Mutations in *REP1* and *REP2* significantly impair segregation fidelity, resulting in failure to transmit plasmid to daughter cells, and subsequent loss from the host population^41^. If copy number drops below a certain threshold within a host cell, 2μ plasmids activate an amplification mechanism that relies on plasmid-encoded *FLP1*^42,43^. *FLP1* encodes a recombinase that creates plasmid structural rearrangements during host S phase via the *FRT* sites, facilitating over-replication via rolling circle replication using host replication machinery^42–46^. *RAF1* encodes a protein that regulates the switch to copy number amplification^47^. Due to this minimal genome, 2μ plasmids rely on host factors for genome replication and segregation during host cell division^44,48–53^

**Figure 1.**
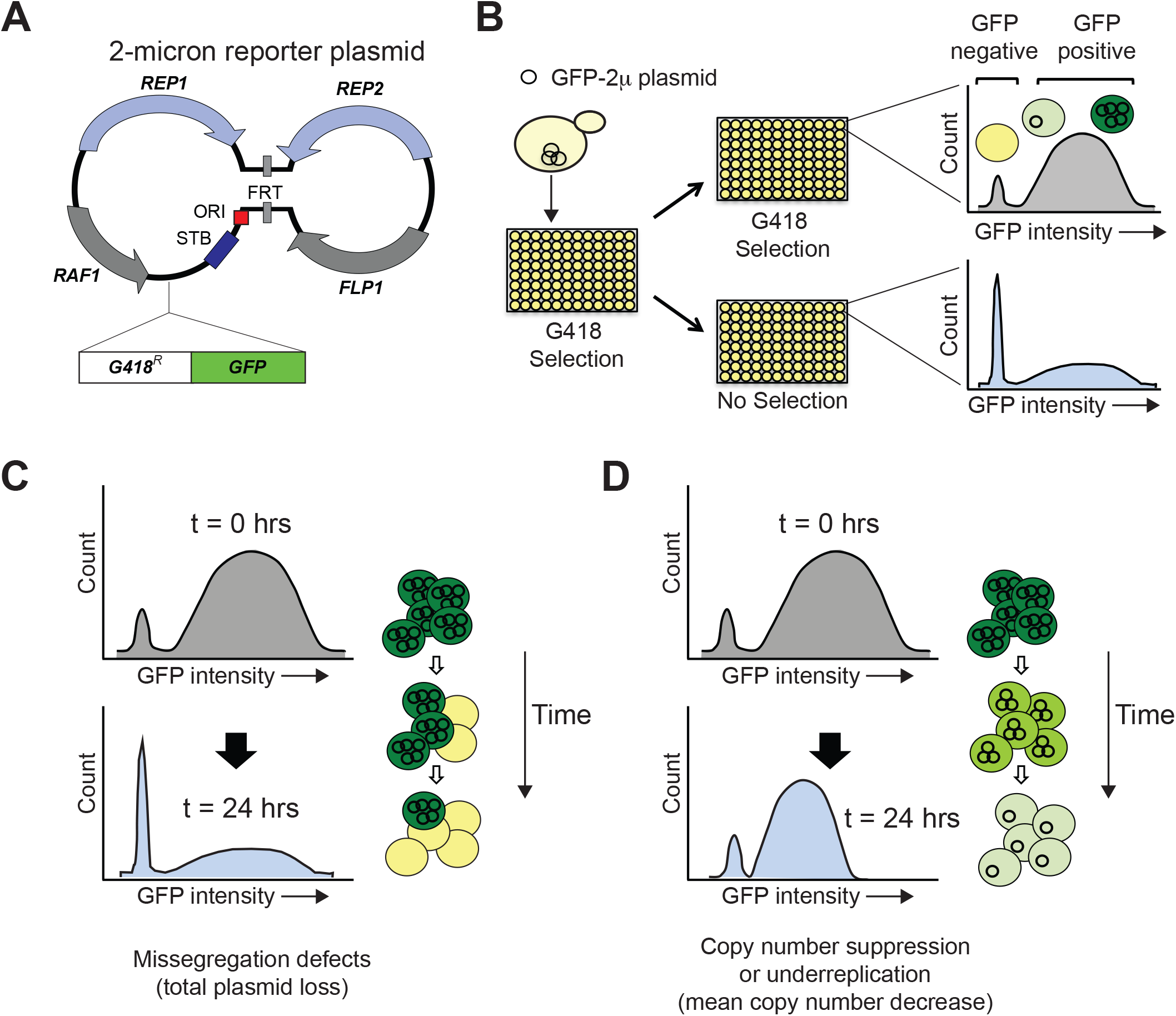
SCAMPR, a novel method to measure 2-micron plasmid stability and dynamics. **(A)** Schematic of GFP-reporter 2-micron plasmid. The endogenous 2-micron encodes an origin of replication (ori), four protein-coding genes (*REP1*, *REP2*, *RAF1*, *FLP1*) and their interacting DNA loci (STB, and FRT). The GFP-2-micron reporter plasmid described here utilizes the full 2-micron genome with an additional integrated G418-resistance and GFP expression cassette. **(B)** A Single Cell Assay for Measuring Plasmid Retention (SCAMPR) utilizes the dual reporter cassette: G418 resistance to ensure plasmid retention while under selection and GFP to facilitate screening of plasmid-positive cells. Cells with the reporter plasmid are kept on G418 selection to ensure the plasmid is present at t=0 and either released to media without selection or passaged with continued G418 selection. Comparing the GFP intensities of the cell populations with and without G418 selection after 24 hours describes the plasmid retention dynamics and population heterogeneity of the host genetic background. SCAMPR can therefore distinguish between alternate mechanisms of plasmid instability, illustrated in (C) and (D). **(C)** Gross segregation defects in which plasmids are not distributed to daughter cells would cause a rapid increase in GFP-negative cells but no change in the median expression of GFP-positive cells. **(D)** Plasmid instability caused by under-replication or copy number suppression would not cause a precipitous decline in GFP-positive cells as in (C) but would instead lead to a reduction in the median GFP intensity of the GFP-positive cells.

Previous studies have identified host factors required by the plasmid. For instance, in addition to DNA replication and origin licensing factors, plasmid require host factors to facilitate proper partitioning into daughter cells, including many spindle-associated proteins ^44,48–52^. Furthermore, host-mediated post-translational SUMO-modification of plasmid-encoded proteins has been shown to have a profound effect on 2μ plasmid stability and host fitness. For example, failure to sumoylate the Rep proteins impairs plasmid stability, whereas deficient sumoylation of Flp1 recombinase leads to recombinase overstabilization, resulting in massively increased plasmid copy number and extreme reduction in host cell fitness^35,54,55^. Indeed, mutations in SUMO E3 ligases *Siz1, Siz2*, the SUMO maturase *Ulp1*, and the SUMO-targeted ubiquitin ligase *Slx8* all lead to hyper-amplification and host cell defects^35,54,55^. These host-plasmid interactions provide potential means for the host to curb deleterious proliferation of plasmids. However, it is unclear whether gain-of-function alleles that restrict or eradicate plasmids exist.

Until recently, 2μ plasmids have been largely omitted from studies of genetic variation in yeast. Although prior work has predominantly focused on canonical A-type 2μ plasmids (found in laboratory *S. cerevisiae* strains), recent studies revealed that 2μ plasmids are quite diverse in budding yeast populations^27,56^. These analyses identified C-type plasmids, extremely diverged D-type plasmids and a 2μ plasmid introgression into *S. cerevisiae* from the closely related species *S. paradoxus*^27,56^. Moreover, previously identified B- and newly described B*-type plasmids were shown to be a result of recombination between A and C types^56–58^. Furthermore, these studies revealed that there are multiple, distinct strains of *S. cerevisiae* that do not harbor any 2μ plasmids. Yet, it remains to be discovered whether 2μ plasmid absence in these strains is the results of stochastic loss or an inherent host trait.

Host cells could influence 2μ plasmid fitness by affecting their copy number, stability, or population heterogeneity. However, these parameters are not captured in traditional plasmid loss assays, which measure either plasmid copy number averaged across the entire population, or plasmid presence versus absence independent of copy number. To quantitatively capture all of these parameters, we developed a new high-throughput, single-cell, plasmid retention assay, SCAMPR (Single-Cell Assay for Measuring Plasmid Retention). We identified three yeast strains that naturally lack 2μ plasmids and reproducibly show a high rate of mitotic instability of 2μ plasmids upon plasmid reintroduction. Focusing on one resistant strain, we used SCAMPR to show that resistance is a dominant, multigenic trait. Using QTL mapping by bulk segregant analysis, we identified one significant genomic locus that impairs 2μ mitotic stability. A candidate gene approach in this locus showed that a single amino acid change in *MMS21* contributes to plasmid instability. *MMS21* is a highly conserved E3 SUMO ligase and an essential component of the Smc5/6 complex, which has not previously been implicated in 2μ biology. Thus, our study reveals a novel pathway by which 2-micron resistance has arisen and persists in natural populations of *S. cerevisiae*.

## Results

### SCAMPR: A Single-Cell Assay for Measuring Plasmid Retention

To determine if there is heritable natural variation in 2μ plasmid stability in *S. cerevisiae* strains, we needed an assay to measure plasmid maintenance at the single-cell level. Traditionally, plasmid loss dynamics have been measured by two types of assays. The first of these is the Minichromosome Maintenance (MCM) assay, in which strains containing plasmids with selectable markers are assessed for plasmid presence versus absence by counting colonies on both selective and non-selective media over time^59^. Due to the labor intensiveness of the assay, MCM is low-throughput since different dilutions need to be tested to recover and reliably measure 30 - 300 colonies per plate. Furthermore, as only a single copy of a selectable marker is required for viable cell growth, substantial variation in plasmid copy number can go undetected by the MCM assay.

A second type of assay traditionally used to measure plasmid stability use molecular methods, such as quantitative PCR (qPCR) or Southern blotting, to assess mean plasmid copy number, relative to genomic DNA, across a population of cells^60^. Compared to the MCM assay, qPCR has the advantages of being high-throughput and not requiring a selectable marker in the plasmid of interest. However, qPCR (or Southern blotting) can only measure the average copy number of a plasmid in a population. Any heterogeneity in plasmid presence or copy number would go undetected by qPCR. Even a combination of the MCM and qPCR assays lacks the resolution to determine the distribution or variability of plasmid copy number within a host population.

We therefore designed a high throughput assay using a reporter 2μ plasmid. To ensure that this plasmid closely resembles endogenous plasmids, we eschewed the use of the yEP multi-copy plasmids commonly used to express yeast ORFs, because they contain only a small portion of the 2μ plasmid. Instead, we built a new 2μ reporter plasmid, which contains both a selectable marker (G418 resistance) as well as a screenable (eGFP) marker, each under a constitutive promoter integrated at a site previously described not to impact plasmid stability^61,62^ (Figure 1A). These dual markers ensured the reporter plasmid could be initially introduced and be retained in plasmid-lacking strains. In addition the GFP allows strains to be easily quantified for plasmid presence, absence, and copy number that scales with GFP intensity in yeast^63–66^. Our analyses revealed that GFP intensity for this 2μ reporter plasmid is roughly normally distributed across single cells (Supplementary Figure S1) indicating that GFP signal did not saturate the detector at high copy number. The endogenous 2μ plasmid loss rates is ~10^−5^ per cell per generation as estimated by colony-hybridization Southern blots^31^. Based on this prior estimate which relied in total loss events, we infer that the stability of the GFP-2μ reporter plasmid is lower than that of the endogenous 2μ plasmid. This could either reflect the difference in precision of plasmid stability measurements or be the result of the constitutive expression of the dual markers. Nevertheless, we conclude that the reporter is well suited for comparative stability studies using the same plasmid in different host backgrounds.

We used this 2μ reporter plasmid with flow cytometry analyses to capture single-cell 2μ fluorescence (Figure 1B), from which we could simultaneously infer both total plasmid loss events by measuring the proportion of GFP-negative cells as well as changes in the median plasmid copy number based on GFP intensity (Figure 1C-D). Importantly, we could also assess the population distribution of GFP intensity, revealing the inherent cellular heterogeneity of plasmid loss and copy number changes. This assay is also higher-throughput than traditional methods. We refer to this assay as SCAMPR (Single-Cell Assay for Measuring Plasmid Retention).

### 2μ plasmid instability in natural yeast isolates is rare and heritable

2μ plasmids are nearly universally present in laboratory strains of *S. cerevisiae*. However, recent studies of natural isolates have revealed a diversity of plasmid types in natural populations, and even strains lacking 2μ plasmids altogether^27,56^. We were particularly interested in plasmid-free strains as these might harbor genetic variants that actively inhibit plasmid stability. To this end, we surveyed a panel of 52 natural *S. cerevisiae* isolates for plasmid presence versus absence via PCR analyses (see Methods). From this panel, we identified three strains (representative gel in Supplementary Figure S2A) that do not contain the 2μ plasmid: Y9 (from ragi, millet), YPS1009 (from oak exudate), and Y12 (from palm wine) (Table 1). To rule out the possibility that PCR surveys were confounded by 2-micron polymorphisms, we also tested these strains via Southern blotting (Supplementary Figure S2B), which supported our conclusion of plasmid absence. Wild diploid strains are homothallic, and capable of mating type switching and self-diploidizing following sporulation. To create stable haploid lines for subsequent analyses, we deleted *HO* endonuclease in the natural isolates before sporulating to produce stable heterothallic haploid strains from each of the three plasmid-free natural yeast isolates (Table 2).

**Table 1.** Natural S. *cerevisiae* isolates screened for the presence or absence of endogenous 2-micron plasmids.

**Table 2.** Engineered *S. cerevisiae* strains used in this study.

Although these three strains lack detectable 2μ plasmids, this absence could be either the result of stochastic loss or host genetic variation that inhibits plasmid stability. Stochastic loss could occur because of rare bottlenecking events in wild populations or during laboratory passaging^15,16^. We predict that such losses would not protect these strains from re-introduction of natural 2-micron plasmids via sex and subsequent propagation^31^. Therefore, if absence were due to stochastic loss, we would expect our reporter 2μ plasmids to be stable in these strains. Alternatively, if the absence of 2μ plasmids reflects true host genetic variation conferring resistance, our reporter 2μ plasmid would be mitotically unstable in these strains. To test these two alternatives, we transformed the GFP-2μ reporter plasmid into haploid cells from these three natural isolates and tested for mitotic plasmid loss using a qualitative colony sectoring assay. As a control, we examined reporter stability in the permissive lab strain BY4742 that was ‘cured’ of its endogenous 2μ plasmid^67^ (see Methods). These analyses showed a clear difference in GFP sectoring (plasmid loss) between the BY4742 laboratory strain and the three natural isolates (Figure 2A).

**Figure 2.**
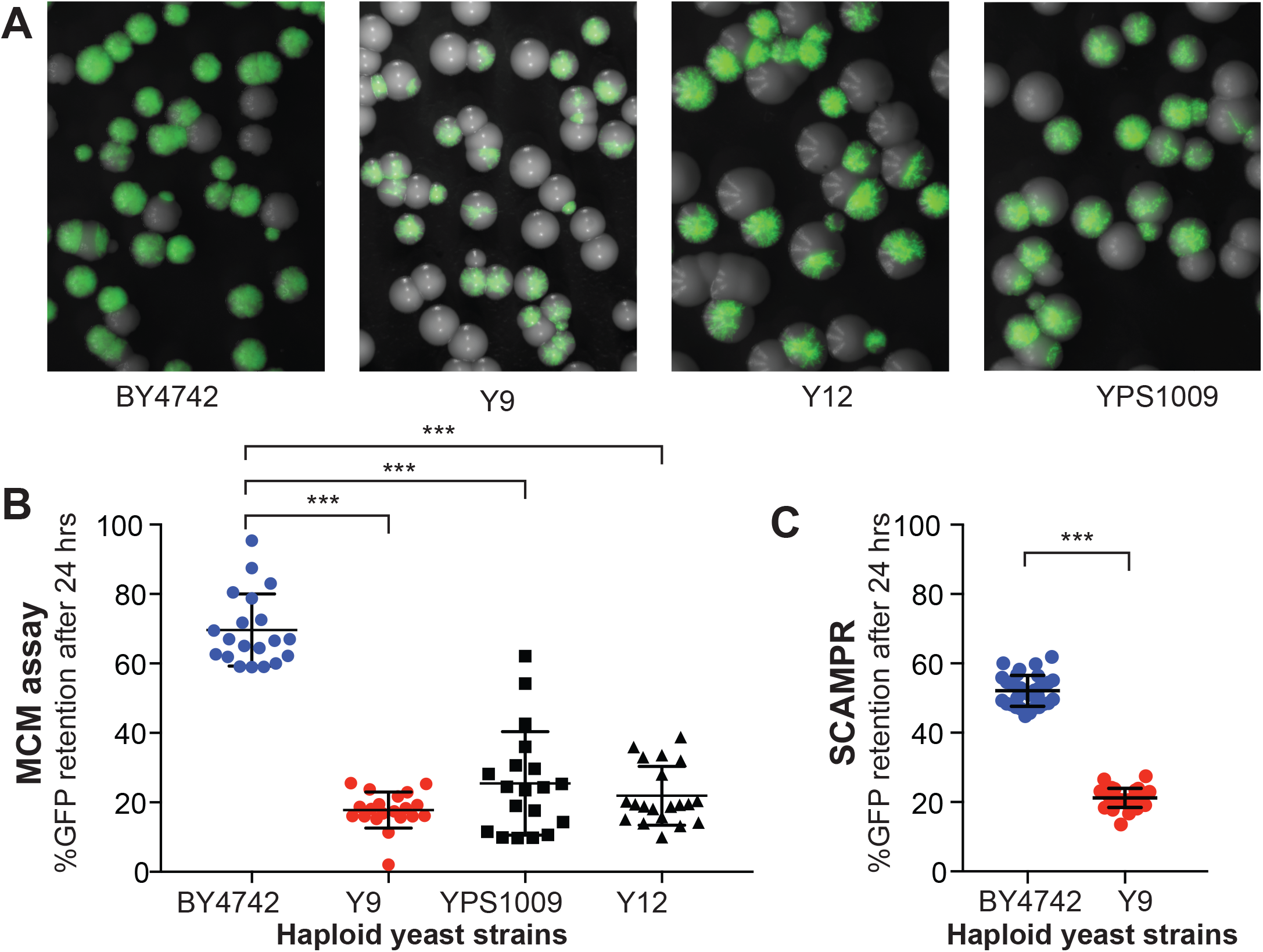
Plasmid instability is a heritable trait in three natural *S. cerevisiae* isolates. **(A)** A colony sectoring assay qualitatively measures GFP-2-micron reporter plasmid loss on solid media. Whereas the majority of colonies in the BY4742 background express GFP, only a small fraction of cells in colonies from wild isolates Y9, Y12, and YPS1009 express GFP. **(B)** The MCM assay quantifies the frequency of 2-micron loss events in different yeast strains. Haploid cells from three wild isolates (Y9, Y12, YPS1009) have significantly lower plasmid retention than haploid cells from the laboratory BY4742 strain. *** p< 0.0001, Kruskal-Wallis test. **(C)** SCAMPR assays confirm that a significantly smaller fraction of Y9 strain haploid cells retain the GFP-2-micron reporter plasmid after 24 hours, relative to haploid BY4742 cells. *** p< 0.0001, Kruskal-Wallis test.

To quantify this difference in plasmid stability between the permissive lab strain and the non-permissive natural isolates, we next measured plasmid stability of the GFP-2μ plasmid over a 24 hour period (~12 generations) using a traditional MCM assay (Figure 2B)^59^. Consistent with the colony sectoring assay, we determined that the reporter GFP-2μ plasmid is significantly less stable in the naturally *cir^0^* wild isolates than in the plasmid-permissive laboratory strain. For example, the Y9 strain maintained 2μ plasmids in only ~5% of cells on average, whereas the BY4742 lab strain maintained plasmids in ~60% of cells. Even after normalization for phenotypic lag (see Methods), we concluded that Y9 and BY4742 strains retain plasmids at 20% versus 70% frequency, respectively. The other two wild strains showed similar plasmid loss frequencies, with the YPS1009 strain exhibiting more variability between replicates than the other two strains. Taken together, these data suggest that 2μ plasmids are mitotically unstable in these three natural isolates. Thus, the absence of endogenous 2μ plasmids in these strains is the result of host genetic variation rather than stochastic plasmid loss.

### Dominant 2μ plasmid instability in the Y9 strain

Of the three natural isolates in which we observed 2μ plasmid instability, the Y9 strain isolate had the least variable plasmid loss phenotype. Furthermore, in a broad analysis of yeast strains, the Y9 strain was found to be phylogenetically close to the Y12 strain^68,69^. Based on this phylogenetic proximity, we hypothesized that Y9 and Y12 strains may share the same genetic basis for host-encoded plasmid instability, which might potentially make this genetic determinant easier to identify. We therefore decided to focus on further understanding the phenotypic and genetic basis of plasmid instability in the Y9 strain.

To infer possible mechanisms of plasmid instability (Figure 1), we further characterized the putative mechanism of 2μ plasmid instability in the Y9 strain using the SCAMPR assay (Figures 2C, S1B-C). We measured the change in distribution of GFP intensity (inferring plasmid copy number changes) among single cells, and total loss events (determined by increase in GFP-negative cells). If the 2μ plasmid were undergoing systematic under-replication due to defects in replication, we might expect an overall and homogenous decrease in median plasmid copy number across the population (Figure 1D). Alternatively, if the 2μ plasmid were being missegregated, we might instead see increasing population heterogeneity, with some cells inheriting no plasmid, while others maintain or even increase plasmid copy number (Figure 1C).

From SCAMPR analyses, we find that even under G418 selection, Y9 cells don’t maintain the reporter 2μ plasmid as well as BY strains (52% versus 90% respectively) (Supplementary Figure S1). Moreover, upon removing pressure to maintain the plasmid (no G418 selection), the proportion of Y9 cells with no GFP (no 2μ plasmid) increases significantly, from 48% to 83%. However, the median GFP-intensity (and inferred 2μ plasmid copy number) of plasmid-bearing Y9 cells remains largely unchanged (Supplementary Figure S1B). We therefore conclude that 2μ plasmid loss in Y9 haploid cells occurs primarily via abrupt, complete loss of plasmids from cells rather than a steady decrease in copy number (Supplementary Figure S1B). This observed pattern of plasmid loss is consistent with plasmid segregation failure during mitosis, rather than a copy number suppression mechanism or plasmid under-replication. As a result of this segregation failure, ‘non-permissive’ Y9 haploid cells lose the 2μ reporter plasmid substantially more quickly than the permissive BY4742 laboratory strain (Figure 2C), mirroring what we initially observed in the colony sectoring assays (Figure 2A).

Next, we investigated whether mitotic instability of the 2μ plasmids in the Y9 strain is genetically recessive or dominant by examining heterozygous diploids of permissive and non-permissive strains. To ensure that mitotic instability was not influenced by ploidy itself, we first tested whether the plasmid instability phenotype we observed in haploid strains persists in homozygous diploid BY4742 and Y9 strains. We found an even bigger difference in plasmid instability between homozygous diploid Y9 and BY4743 strains than between haploid strains (Figures 3A, 2B). We generated a heterozygous diploid strain by crossing the GFP-2μ plasmid containing permissive BY4742 lab strain to the non-permissive Y9 haploid strain. If plasmid loss in Y9 cells were due to inactivating mutations within a host ‘permissivity’ factor required for 2μ mitotic segregation, we might expect plasmid instability to be recessive, with the BY4742 allele providing rescue in the heterozygote. Alternatively, if plasmid instability in Y9 cells were due to a host-encoded, gain-of-function ‘restriction’ factor that impairs mitotic stability of 2μ plasmids, we would expect mitotic instability of 2μ plasmids to be dominant, *i.e.,* heterozygous diploids would also exhibit plasmid instability. We found that heterozygous BY4742/Y9 diploid cells rapidly loses the plasmid after G418 selection is removed (Figure 3A). These findings could result from haploinsufficiency of a permissivity factor, or dominance of plasmid restriction factors in the Y9 genome. We, therefore, considered both possibilities in subsequent analyses.

**Figure 3.**
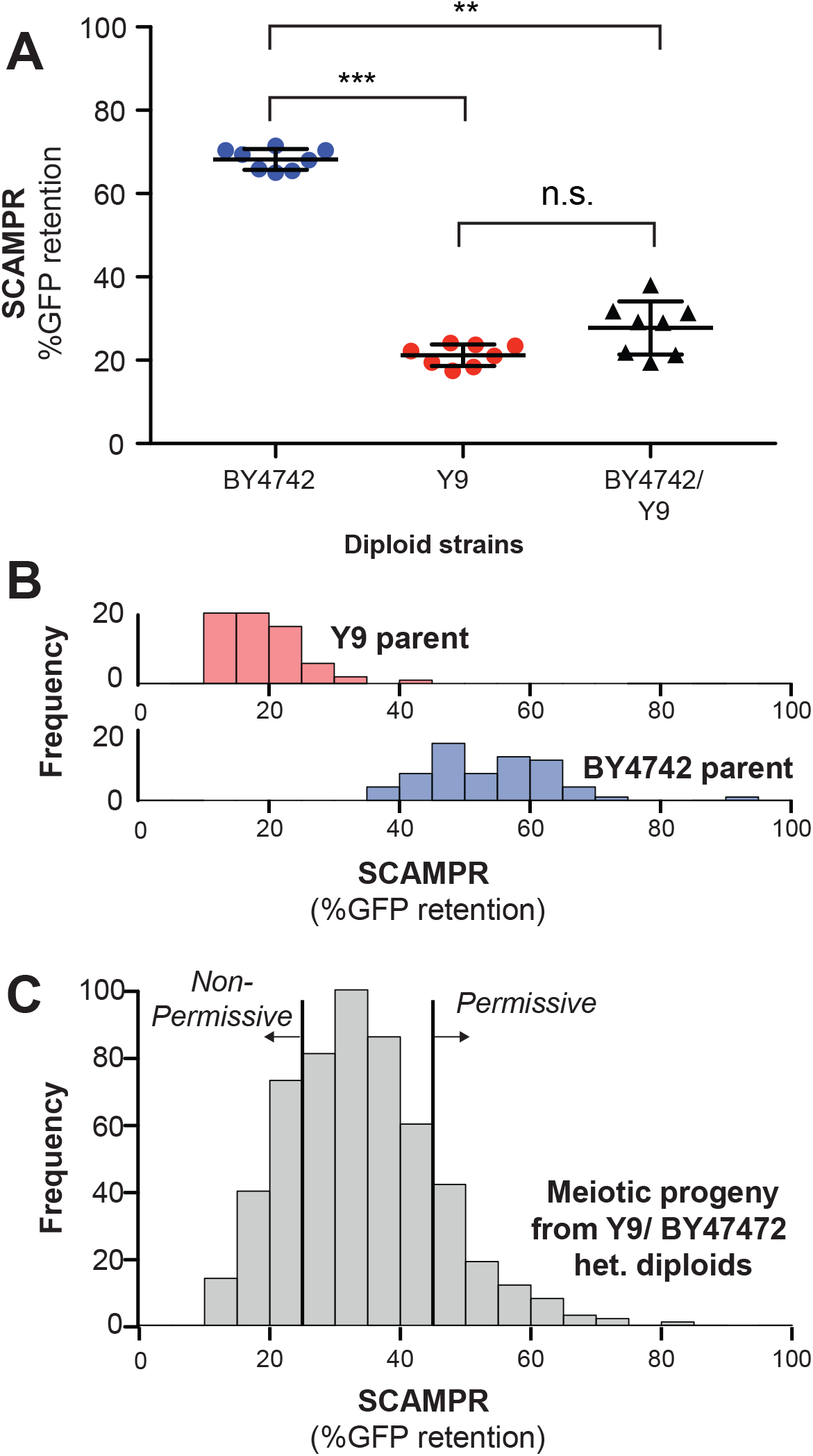
Genetic architecture and dominance of the Y9 plasmid instability phenotype. **(A)** Compared to homozygous BY4742 diploids, heterozygous BY4742/Y9 diploid cells display low plasmid retention after 24 hours, similar to homozygous Y9 diploids. This suggests that the plasmid instability of Y9 cells is a dominant trait. All strains were analyzed with the SCAMPR plasmid retention assay. **p<0.001, *** p< 0.0001, Kruskal-Wallis test; n.s. = not significant. **(B-C)** Progeny phenotype distribution across ~600 random spores (C) shows that most progeny have an intermediate phenotype between the ‘non-permissive’ Y9 parent and the ‘permissive’ BY4742 parent (B). All strains were analyzed in triplicate with the SCAMPR assay. We use this distribution of spores to create cutoffs for the bottom 20% (“non-permissive”) and top 20% (“permissive”) genetic backgrounds for bulk sequencing and segregant analysis.

### Genetic architecture underlying 2μ plasmid instability in the Y9 strain

Previous studies have shown that 2μ plasmids efficiently propagate during meiosis in laboratory strains of *S. cerevisiae*^33,51,70,71^. This propagation is non-Mendelian; even if only one haploid parent initially has 2μ plasmids, all four meiotic progeny typically receive plasmids during meiosis. Because BY4742/Y9 heterozygous diploids exhibit dominant plasmid loss, we maintained G418 selection up to and during sporulation to enrich for tetrads in which all four spores retained 2μ reporter plasmids. We, then, measured plasmid instability phenotypes among meiotic progeny of BY4742/Y9 heterozygous diploids to understand the genetic architecture underlying the Y9 strain’s plasmid instability.

If a single genetic locus were responsible for 2μ plasmid instability, we would expect tetrads to exhibit a 2:2 segregation pattern, with half of the spores phenotypically resembling the permissive BY4742 parent and the other half resembling the non-permissive Y9 parent. Of the 60 tetrads examined, approximately 20% of 4-spore tetrads exhibited a roughly 2:2 segregation pattern, and the remaining 80% tetrads exhibited more complex patterns of inheritance (Supplementary Figure S3). Our results indicate that plasmid instability is heritable but not monogenic. Based on these findings, we used the Castle-Wright estimator^72,73^ to estimate that 2μ plasmid instability is encoded by at least 2-3 independently segregating large effect loci in the Y9 genome.

Next, we performed quantitative trait locus (QTL) mapping using bulk segregant analysis (BSA) to identify the genetic loci that give rise to the Y9 strain’s 2μ plasmid mitotic instability phenotype (Supplementary Figure S4)^74,75^. We selected 600 random spores resulting from a heterozygous BY4742/Y9 diploid containing our reporter 2μ plasmid and used SCAMPR to phenotype plasmid stability^76^. This analysis found that most progeny exhibit intermediate 2μ plasmid stability between BY4742 and Y9 (Figure 3B-C). We then pooled and bulk-sequenced 132 ‘non-permissive’ progeny strains that represented ~20% of progeny with the lowest 2μ plasmid stability and 126 ‘permissive’ strains that represented the ~20% of progeny with the highest 2μ plasmid stability (Figure 3C).

In addition to the progeny pools, we also sequenced the genomes of the three 2μ plasmid-negative strains we identified (Y9, Y12, and YPS1009), as well as UC5, a 2μ plasmid-containing strain (by PCR) that is closely related to Y9 and Y12^68,77^. We mapped reads from these strains back to the *S. cerevisiae* reference genome and created *de novo* assemblies for each strain (see Methods). Unexpectedly, whole genome sequencing revealed that the haploid Y9 parent strain was disomic for chromosome XIV (Supplementary Figure S5A), with the aneuploid chromosome segregating in the Y9 x BY4742 cross. The homothallic Y9 diploid was euploid for chromosome XIV by qPCR and shows a similar plasmid loss phenotype as a homozygous diploid Y9 that has an additional chromosome XIV (Supplementary Figure S5B). These data demonstrate that the aneuploidy for chromosome XIV is not a large contributor to Y9 plasmid instability phenotype^78^. We therefore disregarded the segregating chromosome XIV disomy in our subsequent analyses.

We identified genomic differences between the Y9 and BY4742 strains (see Methods), then compared allele frequencies between the ‘permissive’ and ‘non-permissive’ pools of meiotic progeny from BY4742/Y9 heterozygotes) (Figure 3C, Supplementary Figure S4A). We identified genomic regions in which inheritance of the Y9 allele is significantly more common in the non-permissive progeny pool than the permissive pool (Figure 4A) using the MULTIPOOL algorithm to generate likelihood-based ‘LOD’ (logarithm of the odds) scores^75,79^. We identify loci that are likely linked to the plasmid stability phenotype; although there are a few genomic regions with moderate LOD scores of ~4 (Figure 4B) the most striking LOD score of 9.996 was seen in a high-confidence QTL on chromosome V. While it is challenging to establish a concrete LOD score threshold above which loci are statistically significant, a score of ~10 is comfortably above genome-wide significance thresholds of 3.1-6.3 established empirically in other studies^80–82^. This locus likely encodes the strongest genetic determinant of plasmid instability in the Y9 genome.

**Figure 4.**
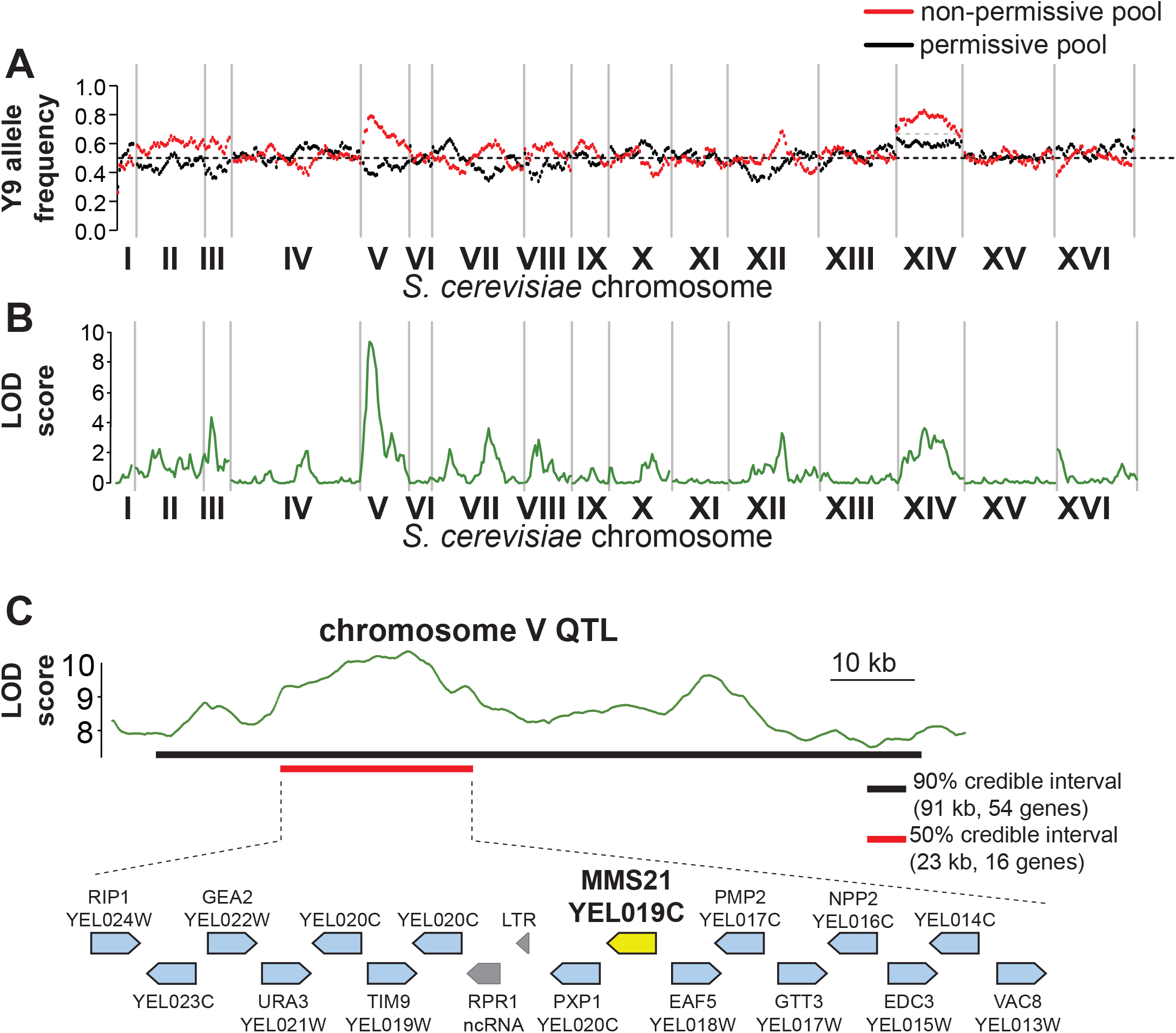
QTL mapping identifies a plasmid instability locus on Y9 chromosome V. **(A)** We plotted the mean Y9 SNP allele frequency in 20kb windows for the ‘non-permissive’ (red) and ‘permissive’ (black) pools of meiotic haploid progeny from BY4742/Y9 heterozygous diploid parents. Associations with a plasmid instability locus would show an increased representation of Y9 alleles in the non-permissive pool and a decreased representation of the BY4742 haplotype in the permissive pool (dotted line indicates equal representation). The increased representation of Y9 alleles on Chromosome XIV in both pools is a result of a segregating disomy in the Y9 parent that we show does not affect the plasmid instability phenotype (Supplementary Fig. S5). **(B)** Based on the allele frequencies shown in (A), we used MULTIPOOL to calculate LOD scores for association with the plasmid instability phenotype. We observe a highly significant LOD score (~10) at a locus on chromosome V. All loci have allele frequencies skewed in the expected direction *i.e.,* the restrictive pool is enriched for Y9 alleles. **(C)** MULTIPOOL 90% (54 genes) and 50% (16 genes) credible intervals for the chromosome V QTL. Among the 16 genes in the 50% credible interval is *MMS21*, which encodes an essential SUMO E3-ligase.

### A single variant of the essential SUMO ligase MMS21 contributes to 2μ mitotic instability in Y9

We focused our efforts on variants within the QTL on chromosome V to identify the genetic basis of Y9-encoded plasmid instability (Supplementary Figure S6). The 90% confidence interval for this QTL is ~91 kb wide and contains 54 ORFs, with a 50% confidence interval 23 kb wide (16 ORFs) (Supplementary Figure S6, Figure 4C). This region contains many polymorphisms between the Y9 and BY4742 genomes but very few structural variants (*i.e.,* large insertions, deletions, translocations). One of the rare large indels is the *URA3* gene, which is present in Y9 and absent in BY4742. Although the *URA3* gene often falls within fitness-related QTLs in BY4742 crosses, detailed follow-up studies (Supplementary Figure S7) allowed us to conclusively rule out a role for *URA3* in the plasmid instability phenotype of the Y9 strain^83,84^.

After excluding dubious ORFs, 44 *bona fide* protein-coding genes remained within the 90% confidence interval. 28 of these candidate genes contained a total of 94 missense changes between Y9 and BY4742, while the rest contained no non-synonymous differences between the parental strains for our QTL cross. We focused on 15 missense polymorphisms (in 11 genes) at which Y9 is identical to the phylogenetically close non-permissive strain, Y12, but different from the closely-related permissive strain, UC5 (Supplementary Table S1). Our attention was drawn to *MMS21*, which contains a single Thr69Ile missense change common to Y9 and Y12, but distinct from the BY4742 laboratory strain and the permissive UC5 strain. Even though *MMS21* has not previously been implicated in 2μ biology, it encodes one of the three mitotic SUMO E3 ligases in *S. cerevisiae*^85^. The two other SUMO E3-ligases, encoded by *SIZ1* and *SIZ2,* have been previously implicated in SUMO-modification of the plasmid-encoded Rep and Flp1 proteins to cause instability or hyper-amplification phenotypes^34,55,86^. Therefore, we evaluated the consequences of Y9’s *MMS21* polymorphism on 2μ plasmid mitotic stability.

We first tested whether Y9 *MMS21* allele was sufficient to confer the plasmid loss trait. We integrated the Y9 *MMS21* allele, with the flanking intergenic regions, into the *ho* locus of BY4742. These engineered BY4742 haploids thus express both the BY4742 and Y9 *MMS21* alleles. Although the addition of the Y9 *MMS21* does lower plasmid stability, this difference is not statistically significant from BY4742 haploids that only express the BY4742 allele (Figure 5A). Thus, the Y9 *MMS21* allele, by itself, does not appear to be sufficient to lower plasmid stability in the BY4742 genetic background.

**Figure 5.**
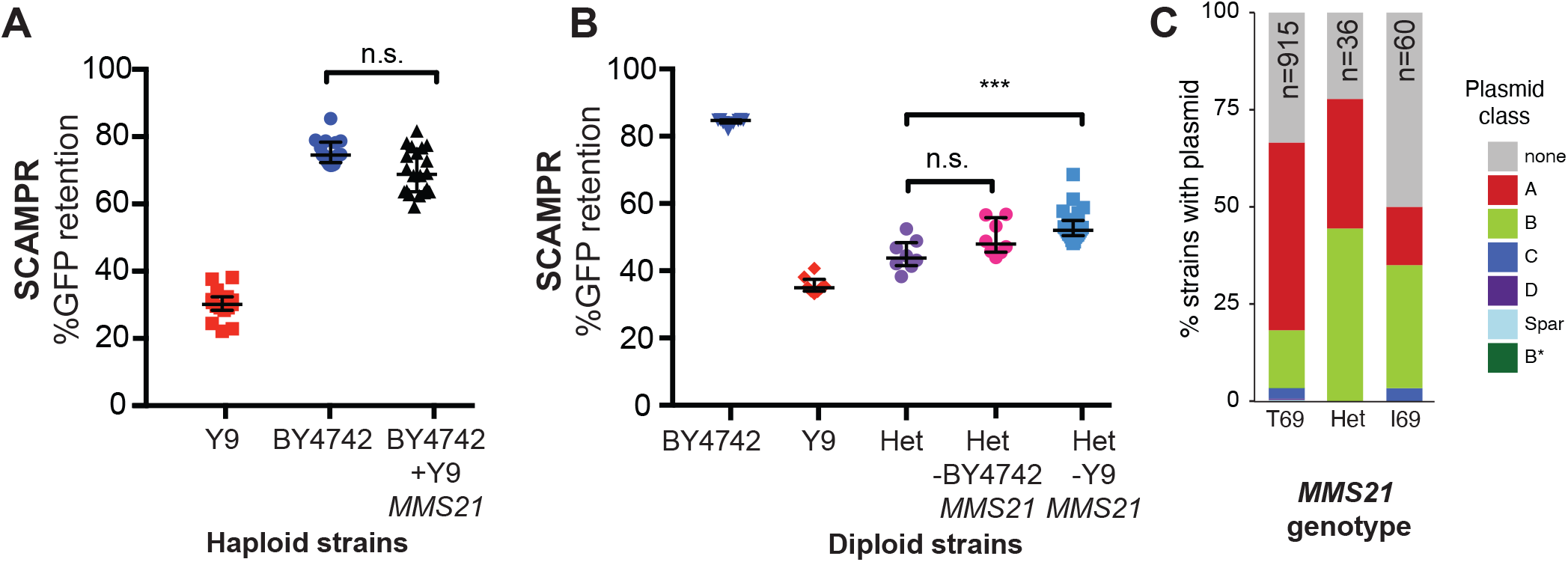
A single SNP in Y9 *MMS21* contributes to the plasmid instability phenotype. **(A)** Introduction of the Y9 *MMS21* allele into BY4742 haploid cells is not sufficient to significantly lower plasmid instability. **(B)** However, removal of the Y9 *MMS21* allele but not the BY4742 *MMS21* allele increases plasmid stability in BY4742/Y9 heterozygous diploids, showing that the Y9 allele of *MMS21* plays an important role in the Y9 plasmid instability phenotype. ** p< 0.001, Kruskal-Wallis test, n.s. = not significant. **(C)** Plasmid prevalence (by plasmid class) for each *MMS21* Thr9Ile genotype within 1,011 sequenced *S. cerevisiae* strains. Plasmid data and genotypes from Peter *et al.*^27^ Strains with the Y9 *MMS21* allele (I69) have a statistically significant decrease in probability of harboring 2-micron plasmids in general, and A-type 2-micron plasmids, in particular. However, this effect can be confounded by the phylogenetic relatedness of these strains.

Next we used reciprocal hemizygosity to test whether loss of Y9 *MMS21* from heterozygous BY4742/Y9 diploids would lead to an increase in plasmid stability. Because *MMS21* is an essential gene, we could not simultaneously delete both the Y9 and BY4742 *MMS21* alleles in heterozygous diploids. Instead, we deleted either the Y9 or the BY4742 allele, yielding BY4742/Y9 diploids that are hemizygous for one or the other *MMS21* allele. If the plasmid stability phenotype were affected by *MMS21* haploinsufficiency, we would expect that deletion of either *MMS21* allele would affect the stability phenotype. Contrary to this expectation, we found that deletion of the Y9 *MMS21* allele, but not the BY4742 allele, results in a reproducible and statistically significant increase in plasmid stability. Loss of the Y9 allele results in a nearly 8% increase in median plasmid stability (Figure 5B). Our results show that the *MMS21* polymorphism clearly contributes significantly to the 2μ mitotic instability of the Y9 strain. Additionally, we conclude that the Y9 plasmid instability trait is caused by dominant plasmid restriction, rather than a haploinsufficient permissivity factor. However, *MMS21* does not explain the entire Y9 phenotype, consistent with plasmid instability segregating as a multigenic trait through the cross. The remaining trait-determining loci in Y9 likely include some of the minor QTL peaks we found but could also include linked polymorphisms within the chromosome V region.

### *MMS21* natural variation within *S. cerevisiae* and between *sensu stricto* species

The Thr69Ile change found in Y9 and Y12 strains is not found in the third non-permissive strain, oak YPS1009, suggesting that YPS1009 acquired 2μ plasmid instability through an independent evolutionary path. The Thr69 allele found in the BY4742 lab strain appears to be the ancestral allele, with Ile69 arising more recently in a subset (96) of 1,011 *S. cerevisiae* strains that were sequenced as part of a recent large-scale study^27^. This study also reported which of the 1,011 strains carry 2μ plasmids. Upon reanalyzing these data, we find that a smaller proportion of *S. cerevisiae* strains homozygous for the *MMS21* Ile69 allele harbor 2μ plasmids compared to strains homozygous for the ancestral Thr69 allele (Figure 5C). In particular, the A-type 2μ plasmids, which we have tested using SCAMPR in this study, appear to be particularly depleted in strains with the Ile69 allele, suggesting that this allele might specifically restrict A-type 2μ plasmids (Figure 5C). While interesting, at present we cannot distinguish whether these observations are a result of a causal association or of shared evolutionary history, due to phylogenetic relatedness of the Ile69 allele-encoding strains.

To explore natural variation in *MMS21* beyond *S. cerevisiae*, we aligned sequences from selected *S. cerevisiae* strains as well as other *Saccharomyces sensu stricto* species and two outgroups (*N. castelli* and *N. dairenensis*) (Supplementary Figure S8). Interestingly, *S. eubayanus* and *S. uvarum* also seem to have independently acquired Ile69 but still harbor endogenous 2μ plasmids, whereas *S. arboricola* has yet another amino acid (alanine) at this position^56^.

The location of the Y9/Y12 amino acid change in the Mms21 protein also provides important clues to its functional consequences. The Thr69Ile change occurs in the third of three alpha-helices in the Mms21 N-terminal domain, which makes contact with the Smc5/6 complex, and is essential for yeast viability^87^ (Supplementary Figure S9). Yeast cells deficient for *MMS21* show gross chromosomal segregation defects and die as large, multi-budded cells^88^. However, the C-terminal zinc finger RING domain responsible for sumoylation of substrates is dispensable for Mms21’s essential function^87^. We therefore speculate that the non-permissive *MMS21* allele may act by directly affecting the Smc5/6 complex rather than through its sumoylation function. Despite the Smc5/6 complex’s essential role in the removal of DNA-mediated linkages to prevent chromosome missegregation and aneuploidy, it has not been directly implicated in 2μ stability. Our finding that a single polymorphism at the Mms21-Smc5 interaction interface reduces 2μ stability thus reveals a novel facet of host control.

## Discussion

In this study, we leveraged natural variation to identify a gain-of-function variant that restricts 2μ plasmids in *S*. *cerevisiae*. Our approach is complementary to the traditional biochemical and genetic approaches that have previously used loss-of-function genetic analyses to study host regulation of 2μ plasmids. Natural variation studies can identify alleles of host genes that retain host function but still block SGEs like 2μ plasmids. Such studies can reveal novel mechanisms of host control, which may be otherwise challenging to discover via loss-of-function analysis.

Although 2μ-based vectors have long been used as an important tool in yeast genetics, studies of 2μ plasmids as natural SGEs have lagged behind considerably. Our new phenotyping assay, SCAMPR, makes the 2μ plasmid a more tractable system. SCAMPR captures single cell data that facilitate studies of population heterogeneity, allowing inferences of the mechanisms by which plasmids may be controlled by their hosts. Although recent advances in single-cell genome sequencing make it possible to directly sequence and infer copy number of 2μ plasmids, this would be prohibitively expensive compared to the GFP-based FACS profiling methodology we use. SCAMPR has potential for expanded use, for example, to explore meiotic plasmid transmission dynamics. SCAMPR could also be paired with host lineage tracking to assess plasmid fitness burden alongside plasmid loss dynamics in competitive fitness assays. This paired strategy would provide a powerful approach for understanding the relative contribution of both plasmid fitness cost and host-plasmid incompatibility across hosts. In general, SCAMPR could be utilized to study high-copy number SGE plasmid dynamics, DNA replication and segregation, in any system where expression is well matched to copy number.

Our survey of 52 wild *S. cerevisiae* isolates identified three strains that naturally lack 2μ plasmids. Detailed studies of one of these strains, Y9, revealed that 2μ plasmid instability is heritable, dominant and likely the result of multiple contributing alleles. Through QTL mapping by bulk segregant analysis, we identified a significant locus on chromosome V associated with 2μ plasmid loss. We found that a single amino acid variant in Y9 *MMS21*, which encodes an essential SUMO E3 ligase in *S. cerevisiae*, contributes to 2μ plasmid instability. *MMS21* does not fully account for the 2μ plasmid loss phenotype in heterozygous BY4742/Y9 strains. This result is unsurprising based on our tetrad analysis and QTL mapping, which both suggest additional independently segregating loci affect plasmid stability. Although loss of Y9 *MMS21* from heterozygous diploids leads to a relatively modest effect on 2μ plasmid instability, it may still account for all of the QTL signal we observe in chromosome V. Alternatively, the QTL on chromosome V may contain additional determinants of plasmid instability in close genetic linkage to *MMS21*. CRISPR-Cas9 based approaches will be useful to test a large number of genomic changes rapidly and in parallel between Y9 and BY4742 to identify other determinants of plasmid instability in this QTL and in other candidate loci^89,90^.

The reciprocal hemizygosity experiment (Figure 5B) reveals that the Y9 variant of *MMS21* acts dominantly to restrict mitotic stability of 2μ plasmids, ruling out the possibility that this allele is haploinsufficient. We considered two possibilities by which this allele may exert its dominant effect on plasmid stability. The first possibility is that the Y9 allele of *MMS21* encodes a dominant-negative allele, which impairs the function of the Smc5/6 complex whether in a haploid or heterozygous state. Although this impairment does not negatively impact essential host functions when 2μ plasmid is absent (Y9 strains are viable and fit), this allele would have a fitness deficit in the presence of 2μ plasmids, as host functions become overburdened when hijacked by the parasite. Under this scenario, impairment of Mms21 function would be synthetically lethal with high levels of 2μ plasmid. If this were the case, we would expect to see decreased viability of Y9 cells upon 2μ reintroduction, or appearance of *nibbled* colony morphology as seen in *ULP1* mutant cells harboring high levels of 2μ plasmids. However, we did not observe such deficits upon reintroduction of 2μ plasmids into Y9 (see Figure 2A for example). Moreover, plasmid-restrictive host alleles would only arise and propagate in natural populations if their fitness cost did not outweigh the modest 1-3% fitness cost imposed by the widespread 2μ plasmids in *S. cerevisiae* populations. We, therefore, favor the second possibility, in which the Y9 *MMS21* allele represents a separation-of-function allele that is still capable of performing host functions but impairs 2μ mitotic stability.

The 2μ plasmids appear to have co-evolved with budding yeasts for millions of years and are prevalent in species such as *S. cerevisiae*. Long-term coevolution appears to have “optimized” 2μ plasmids as tolerable parasites: not too great of a burden on host fitness, but still high enough plasmid copy numbers to ensure stable propagation. This copy number balance is achieved through both plasmid (*e.g.,* Flp1 repression) and host (e.g., sumoylation) contributions. Nevertheless, there are hints that this truce between 2μ plasmids and yeast may be uneasy. 2μ plasmid stability is frequently compromised in heterospecific (other species) hosts, suggesting it is actively adapting to maintain stability within its native budding yeast species^37^. Our discovery of a natural host variant of *S. cerevisiae* that impairs conspecific (from same species) 2μ plasmid stability further supports the hypothesis that even low fitness costs invoked by 2μ plasmids are sufficient to elicit a host evolutionary resistance response.

Most studies of 2μ plasmid plasmids (including this one) have focused on the A-type variant that is most commonly found in laboratory strains. However, new sequencing studies have revealed that *S. cerevisiae* strains harbor a diverse set of 2μ plasmid plasmids^27,56^. This diversity of 2μ plasmid plasmids might itself have arisen as a result of host defenses within *S. cerevisiae,* leading to plasmid diversification. For instance, although SCAMPR studies revealed the importance of the Y9 *MMS21* variant against the stability of the A-type plasmid, it is possible that this variant is ineffective against the B-type variant. Thus, 2μ plasmids might exist in a frequency-dependent regime with their budding yeast hosts; A-type plasmids might thrive in certain host genetic backgrounds whereas B-type plasmids might thrive in others. The simultaneous presence of multiple 2μ plasmid types within species could explain the presence of standing variation in plasmid instability phenotypes in *S. cerevisiae* populations, including the low observed frequency of the Y9 *MMS21* allele.

Testing the effects of *MMS21* and other restrictive alleles on stability of different 2μ plasmids (e.g. B- or C-type) might provide a means to distinguish between universal versus plasmid-type-specific restriction. Future studies could employ SCAMPR to study the functional consequences of the natural diversity of 2μ plasmids in yeast. In particular, SCAMPR reporters from different types of *S. cerevisiae* 2μ plasmids and from divergent *Saccharomyces* species may reveal important biological determinants behind their co-evolution and long-term success in budding yeast species.

The identification of *MMS21* led us to initially suspect that this locus might represent another connection between the SUMO-ligation machinery and 2-micron plasmid stability^34,35,55,91^. However, the location of the Thr69Ile missense change at the binding interface between Mms21 and Smc5 (Supplementary Figure S9) suggested an alternate mechanism that relies on the Smc5/6 complex. The Smc5/6 complex lies at the nuclear periphery, where it anchors dsDNA breaks to facilitate repair, resolves X-shaped DNA structures that arise during DNA replication and repair, and helps mediate sister chromatid cohesion^88^. All three of these cellular processes directly impact stability of 2μ plasmids, which also physically localize to the nuclear periphery^92^. Interference with Mms21, an essential component of the Smc5/6 complex, could thus directly affect both segregation of 2μ plasmids as well as interfere with their amplification via Flp1-induced recombination intermediates. We therefore speculate that the Y9 Mms21 variant may restrict 2μ stability through Smc5/6, rather than through sumoylation.

Although Smc5/6 has not previously implicated in 2-micron stability, this complex has been previously implicated in the stability of viral episomes. Indeed, human Smc5/6 acts as a restriction factor against hepadnaviruses such as human Hepatitis B virus^93,94^. To counteract this restriction function, diverse hepadnaviruses encode antagonist HBx proteins that degrade mammalian Smc5/6 and restore viral fitness^93–95^. Our findings that components of the yeast Smc5/6 complex affect 2μ stability suggest that the Smc5/6 complex might provide a general mechanism to protect host genomes from the deleterious consequences of multicopy genetic parasites.

## Methods

### Strain growth and construction

For most experiments, yeast strains were grown in standard yeast media at 30°C unless otherwise noted^76^. Plasmid transformation was carried out using a high efficiency lithium acetate method^76^. The GFP-2-micron plasmid was created by Gibson assembly directly into otherwise plasmid-less yeast strains *cir^0^* BY4741 (*MAT****a*** haploid) and BY4742 (*MAT****α*** haploid), which had been cured of their endogenous plasmids by previously published methods^67,96^. To avoid disruption of the plasmid’s endogenous replication and segregation machinery, a cassette containing both markers was integrated into the A-type 2-micron sequence found in the *S. cerevisiae* laboratory strain BY4741 at a restriction site reported to tolerate insertions of up to 3.9 kb without impacting copy number or stability^62^. We did not use any bacterial cloning vector sequences to minimize unnecessary or destabilizing changes to the 2-micron reporter plasmid, so the reporter plasmid was directly assembled in yeast. Assembling in *cir^0^* strain backgrounds avoided multiple plasmid genotypes within a strain background that could have led to plasmid competition or recombination.

We used the NEBuilder HiFi DNA Assembly Master Mix (product E2621) for Gibson assemblies. Yeast plasmids were recovered using Zymoresearch Zymoprep Yeast miniprep kit (D2004). The assembled plasmids were then retransformed to the same *cir0* yeast backgrounds to ensure plasmid clonality. Genetic crosses were carried out on a Singer Sporeplay dissection scope, for both tetrad dissection and selection of unique zygotes for mating strains. Strain mating type was confirmed by halo formation in the presence of known mating type tester strains (see strain table). Strains used in this work are listed in Table 1.

Natural isolates were obtained as homothallic diploids (capable of mating type switching and self-diploidization). We made stable heterothallic haploid strains (no longer capable of mating type switching) by first knocking out *ho* endonuclease prior to sporulation (*hoΔ::HphNT1*). We found that the natural isolates required significantly longer homology arms for proper DNA targeting when making integrated genomic changes (e.g. gene deletions) via homologous recombination. Where BY4742 lab strains utilized ~50bp homology arms for high efficiency recombination, Y9 required ~1kb flanking homology. Even with longer homology, a substantial number of clones in any transformation did not contain the desired edit. These hurdles made editing the Y9 genome challenging.

### Colony sectoring

Confirmed transformants were cultured under G418 selection, then plated to YPD medium where colonies were allowed to form without selective pressure to maintain the reporter plasmid. After 2 days growth at 30°C, colonies were imaged under white light and GFP excitation to assess qualitative plasmid loss in the different strains using a Leica M165 FC dissection scope with a GFP filter and Leica DFC7000 T camera. Colony sectoring was visually assessed. We then performed image processing using ImageJ to split channels and recolor the GFP channel.

### MCM assay

MCM assays were performed as previously published^59^. However, samples were taken at only two time points. Therefore, we reported changes in frequency of 2-micron plasmid rather than an estimated rate of loss per generation. This two-timepoint measurement also provided a more direct comparison to the SCAMPR assay. At time = 0 hours and 24 hours, cells are plated on both selective and non-selective media to determine what fraction of the population maintains the plasmid by virtue of encoding the selectable marker. Samples were plated at multiple dilutions to ensure between 30-300 CFU per plate. All strains containing the reporter plasmid were grown under G418 selection to ensure 2-micron plasmid presence prior to the start of the assay. At time = 0 hours, cultures were transferred into liquid media with shaking, but without drug selection for 24 hours. After 24 hours, cultures were diluted in PBS and plated on YPD either with or without G418 selection at multiple dilutions, targeting 30-300 CFU per plate. Plates were incubated for 2 days, then colonies were manually counted to determine what fraction of the population were G418 positive. Calculations were based on whichever dilution gave a countable (30-300 CFU) plate. Multiple replicates (at least 8) were done for each strain to measure variability in plasmid retention. A subpopulation of GFP negative, G418-negative cells can be found even under selection. This ‘phenotypic lag’ occurs because of protein persistence following DNA loss. Additionally, while cells that lose the plasmid die under selection, other plasmid-free cells are constantly generated as well. We, therefore, normalize data to account for different starting frequencies of plasmid negative cells by comparing cells grown with or without G418 selection for 24 hours (Figure 1B).

### SCAMPR

SCAMPR samples were prepared as for MCM assay. When grown in 96-well format at 30°C, cultures were shaken using a Union Scientific VibraTranslator to ensure aeration. Fluorescence was directly measured by flow cytometry at 0 and 24 hour timepoints. A BD Canto-2 cytometer was used to collect cell data. FlowJo software was used for subsequent data analysis: samples were gated for single cells, omitting doublets/multiple cell clumps and any cell debris. Single cells were gated for GFP-positive and -negative populations, using GFP negative strains and single-copy integrated GFP-positive strains as gating controls. Summary statistics (frequency of GFP-positive and -negative cells, GFP intensity) were exported from FloJo. Each strain was measured in at least triplicate per assay and means are reported here.

### Statistical analyses

For SCAMPR and MCM assay results, we determined significance by non-parametric tests. In the case of comparing two strains we used two-tailed Mann-Whitney, or for comparing three or more strains, Kruskal-Wallis with Dunn’s multiple comparison test. Graphs were prepared and statistical analysis done using GraphPad Prism 7 software.

### Screening for endogenous 2-micron plasmid in natural isolates of S. cerevisiae

Natural isolates (Table 1) were generously shared by Dr. Justin Fay. DNA from these strains was isolated using standard Hoffman and Winston preps, then probed by PCR and Southern blot^76^. Two pairs of primers were designed to amplify either *REP1* or *FLP1* (FLP1_F: CCACAATTTGGTATATTATG, FLP1_R: CTTTCACCCTCACTTAG, REP1_F: AATGGCGAGAGACT, REP1_R: CGTGAGAATGAATTTAGTA), the two best conserved coding regions of the plasmid as previously described^57^. Only strains that showed negative PCR results for both sets of primers were further validated by chemiluminescent Southern blot using the Thermo North2South kit (17097). Briefly, whole genome DNA was digested, run on an agarose gel in TAE, transferred to membrane and probed with chemiluminescent probes created from digested endogenous 2-micron plasmid collected from BY4741 by Zymoresearch yeast plasmid miniprep kit (D2004).

### Illumina sequencing, library preparation, and QTL mapping via bulk segregant analysis

We prepared high quality genomic DNA for sequencing using Zymoresearch Yeastar kits with per manufacturers instructions (D2002 - using chloroform method). Sequencing libraries using the TruSeq method for genomic DNA (Illumina). Samples were multiplexed and run on an Illumina HiSeq by the Fred Hutchinson Sequencing core facility to generate 50bp paired-end sequences (SRA accession PRJNA637093). 100bp paired-end reads for the lab strain, BY4742, were downloaded from the SRA database (accession SRR1569895). Reads that failed Illumina’s ‘chastity filter’ were removed using a custom R script, and adapters and low-quality regions were trimmed using cutadapt with parameters -q 10 --minimum-length 20^97^. Trimmed read pairs were aligned to the sacCer3 reference genome assembly using BWA-backtrack^98^. Mean coverage in non-overlapping 20kb windows across the genome was calculated and plotted using R and Bioconductor.

For bulk segregant analysis, we first identified a conservative set of 47,173 high quality SNPs that distinguish the cross parents (Y9 and BY4742) as follows. Before SNP-calling, BWA output files were processed using Picard’s MarkDuplicates tool and indels were realigned using GATK’s RealignerTargetCreator and IndelRealigner tools^99,100^. We then called SNPs using samtools mpileup (parameters --skip-indels -t DP -uBg -d 6660) and bcftools call (parameters -vmO z -o), and counted reads matching each allele using GATK’s VariantAnnotator DepthPerAlleleBySample module (with --downsampling_type NONE option)^101^. We used R and Bioconductor to further filter SNPs to obtain the final set of 47,173 SNPs, removing any that overlapped repetitive elements, SNPs with QUAL score <200, SNPs with unusual coverage in any sample, and SNPs with an apparent mix of alleles in either of the haploid parental strains. We then ran MULTIPOOL in ‘contrast’ mode on allele frequencies at each SNP in the permissive and non-permissive pools to generate LOD scores across all chromosomes^79^.

To identify candidate functional polymorphisms, we took two approaches: (a) we performed more sensitive SNP-calling, including small insertions and deletions; (b) to detect larger insertion/deletion events, we generated *de novo* assemblies from each strain, aligned them to the reference genome assembly, and identified locations where assemblies differed. In more detail, the first approach used processed alignments (see above) as input to GATK’s HaplotypeCaller (parameters -stand_call_conf 30.0 - stand_emit_conf 10.0)^100^. Functional consequences of each variant were annotated using Ensembl’s Variant Effect Predictor^102^. For the second approach (*de novo* assemblies), we performed error correction on the adapter-trimmed reads using musket (parameters -k 28 536870912) and then used SOAPdenovo2 across a range of k-mer sizes and fragment sizes, choosing the combination for each sample that yielded the assembly with highest N50 length as determined using QUAST (Genbank accession numbers forthcoming)^103–105^. We obtained tiling path alignments of each assembly to the sacCer3 reference genome assembly using MUMMER (nucmer parameters -maxmatch-l 100 -c 500, delta-filter options -m)^106^. Structural variants were determined from genome alignments using Assemblytics (variant size range 1bp-100kb)^107^.

### Structure visualization

The Cn3D viewer was used to visualize Thr69Ile on a crystal structure of *MMS21* with *SMC5* made available by Duan *et al*^87,108^.

### Analysis of MMS21 natural variation

To examine natural variation in *MMS21* across *S. cerevisiae* strains and in other fungal species, we first extracted the *MMS21* (YEL019C) open reading frame from the reference assembly (sacCer3, chrV:120,498-121,301, - strand) and translated that sequence. We then used this *MMS21* protein sequence as the query in tblastn searches against various databases^109^. Searching the NR database, using taxonomic restrictions as needed, yielded *MMS21* sequences from *S. paradoxus* (XM_033909904.1), *S. eubayanus* (XM_018364578.1), *S. jurei* (LT986468.1, bases 125,344-126,147, - strand), *S. kudriavzevii* (LR215939.1, bases 100238-101041, - strand), *N. castellii* (XM_003677586.1) and *N. dairenensis* (XM_003671024.2). For additional orthologs, we downloaded individual genome assemblies from NCBI and performed local tblastn searches for *S. arboricola* (GCA_000292725.1; *MMS21* at CM001567.1:97,893-98,699, - strand), *S. mikatae* (GCA_000166975.1; *MMS21* at AABZ01000034.1:31,665-32,468, - strand), *S. uvarum* (GCA_002242645, *MMS21* at NOWY01000012.1:107,550-108,353, + strand). Additional *S. cerevisiae* strain sequences come from our own *de novo* assemblies, where we used blastn to identify *MMS21*.

*MMS21* genotypes in the 1,011 isolates previously sequenced^27^ were accessed via that publication’s supplementary data file 1011Matrix.gvcf.gz. Plasmid status was obtained from another supplementary file (Table S1) and cross-referenced with genotype in R.

## Acknowledgements

We thank past and present members of the Malik, Dudley and Raghuraman/Brewer labs for helpful discussions throughout this project. We especially thank Aimée Dudley, Mosur Raghuraman, Gavin Sherlock, Tera Levin, Antoine Molaro, Courtney Schroeder, and Yu-Ying (Phoebe) Hsieh, for their comments on the manuscript. We are grateful to Justin Fay for the generous gift of natural yeast isolates. We appreciate all the generous assistance and advice from Flow Cytometry and Genomics shared resource facilities at the Fred Hutchinson Cancer Research Center. This work was supported by an NSF graduate research fellowship (Grant No. DGE-1256082 to M.H.), NIH/NHGRI Genome Training Grant at the University of Washington (5T32HG000035-20 to M.H.), NIH R01 grant GM074108 (to H.S.M.) and an Investigator award from HHMI (to H.S.M.). The funders had no role in study design, data collection and analysis, decision to publish, or preparation of the manuscript.

## Supplementary Materials

**Table S1. Missense polymorphisms in the 90% credible QTL interval for plasmid instability** Missense polymorphisms shared between the non-permissive Y9 and Y12 strains, but distinct from the closely-related plasmid-permissive strain, UC5, and the permissive laboratory strain, BY4742, are listed.

**Figure S1.** SCAMPR analysis for permissive and non-permissive *S. cerevisiae* strains. **(A)** SCAMPR analysis in the laboratory BY4742 strain reveals that GFP intensity for the 2-micron reporter plasmid is roughly normally distributed across single cells. Upon relaxation of G418 selection, there is an increase in the number of cells lacking 2-micron plasmid from ~10 to 24%, although the median GFP intensity of plasmid-bearing cells remains mostly unchanged. **(B)** In non-permissive Y9 strains, there is an increase of plasmid -lacking cells from 48% to 83% upon relaxation of G418 selection. Again, the median GFP intensity of plasmid-bearing cells remains largely unchanged. From these analyses, we conclude that plasmid instability in Y9 cells occurs via mis-segregation defects. **(C)** Table summarizing the plasmid-negative cell fractions in BY4742 and Y9 cells, with and without G418 selection.

**Figure S2.** Three natural *S. cerevisiae* isolates lack endogenous 2-micron plasmids. **(A)** Representative PCR analysis shows that most of the 52 natural isolates tested harbor endogenous 2-micron plasmids, except for three strains (one indicated). Drosophila DNA was included as a negative control template. Representative gel shown for the presence of *REP1* (Methods) **(B)** Southern blot analysis confirms the absence of endogenous 2-micron plasmids in the 3 natural isolates Y9, Y12, and YPS1009 as compared to the BY4742 positive control.

**Figure S3.** Tetrads dissected from meiosis of B4742/Y9 heterozygous diploids reveal a range of plasmid stability phenotypes. All four spores dissected from meiotic tetrads were individually assayed by SCAMPR. While some tetrads display a 2:2 segregation pattern of plasmid instability/stability consistent with a single Mendelian locus, others suggest a more complex inheritance pattern. This pattern is consistent with at least 2-3 independently segregating loci in the Y9 genome that inhibit 2-micron plasmid stability. Values plotted are the mean of SCAMPR measurements of three replicates per progeny.

**Figure S4.** Schematic of QTL mapping by bulk segregant analysis^110^. We crossed non-permissive Y9 and permissive BY4742 haploid cells to create heterozygous diploids. We expect any Y9 alleles associated with plasmid instability (yellow star) to be concentrated in pools of meiotic progeny that show plasmid instability with SCAMPR analyses. Therefore, by determining where the Y9 allele frequency is elevated in the non-permissive pool and depleted in the permissive pool, we can highlight genetic loci that are likely to significantly contribute to the plasmid instability phenotype. We use allele frequencies as input to the MULTIPOOL algorithm to calculate a LOD scores to indicate statistical likelihood of each genetic locus contributing to the plasmid instability phenotype^79^.

**Figure S5.** Aneuploidy of chromosome XIV in Y9 strain. **(A)** Whole genome sequencing reveals ~2X coverage of chromosome XIV in the sequenced Y9 haploid strain, indicating that this parent is disomic for this chromosome. This disomy segregates among the meiotic progeny from BY4742/Y9 heterozygous diploids. **(B)** Y9-derived diploid strains that were euploid or aneuploid for chromosome XIV were identified by qPCR and phenotyped^78^. We find no difference in plasmid instability phenotypes, indicating that disomy of chromosome XIV is not likely to contribute to this trait.

**Figure S6. Y9 Chromosome V is most strongly associated with plasmid instability. (A)** Over-representation of Chromosome V Y9 alleles in the non-permissive pool (red) and under-representation in the permissive pool (black). **(B)** Based on the allele frequencies shown in (A), we calculated LOD scores and 50%, 90% credible intervals for association with the plasmid instability phenotype.

**Figure S7.** Deletion of URA3 from Y9 haploid cells does not affect their plasmid instability phenotype. Independently derived biological replicates (Rep1 through 5) of Δura3 Y9 cells are not different from wildtype Y9 in terms of their plasmid instability phenotypes. *** p< 0.0001, Kruskal-Wallis test, n.s. = not significant.

**Figure S8**. Sequence of *MMS21* codon 69 across the *Saccharomyces sensu stricto* clade, as well as selected *S. cerevisiae* strains and two outgroup *Naumovozyma* species. Species cladogram was adapted from previous studies^111,112^. This analysis shows that the Thr69 allele of *MMS21* is ancestral in *S. cerevisiae* but is not universally conserved in closely-related species.

**Figure S9.** The Thr69Ile polymorphism is located at the Mms21-Smc5 binding interface. **(A)** The Thr69Ile polymorphism occurs in the third alpha-helix in the Mms21 N-terminal domain. The C-terminal domain of *MMS21* encodes the SUMO E3-ligase associated RING domain. **(B)** Schematic of co-crystal structure of Mms21 with the coiled coil domain of Smc5 (PDB: 3HTK) shows that the Thr69 residue in the N-terminal domain is at the direct binding interface between the two proteins^87^. **(C)** Cartoon of Smc5/6 complex in *S. cerevisiae* showing the Mms21-Smc5 interaction is adapted from a previous review^113^.

