## Supplementary Figures S1-S9 for "A natural variant of the essential host gene *MMS21* restricts the parasitic 2-micron plasmid in *Saccharomyces cerevisiae*"

A

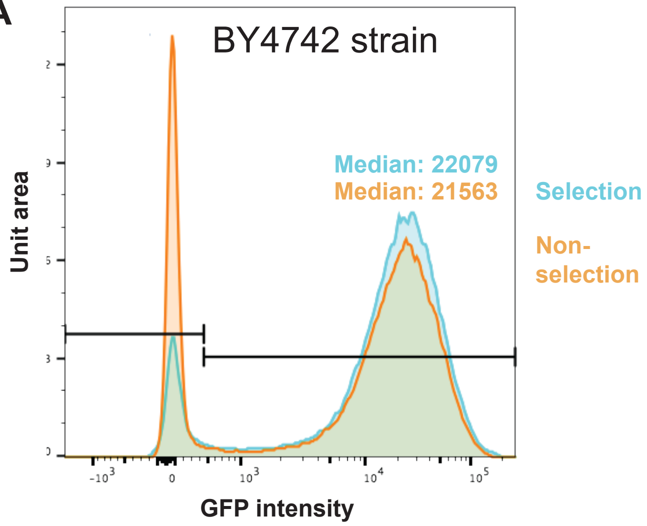


B

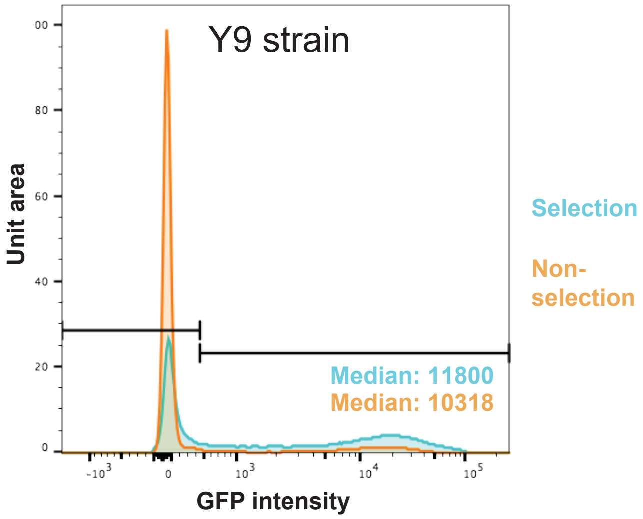


C

| | Selection (Plasmid-negative %) | Non-Selection (Plasmid-negative %) |
|---|---|---|
| BY4742 | 9.7 | 24 |
| Y9 | 48 | 83.3 |

### Supplementary Figure S1

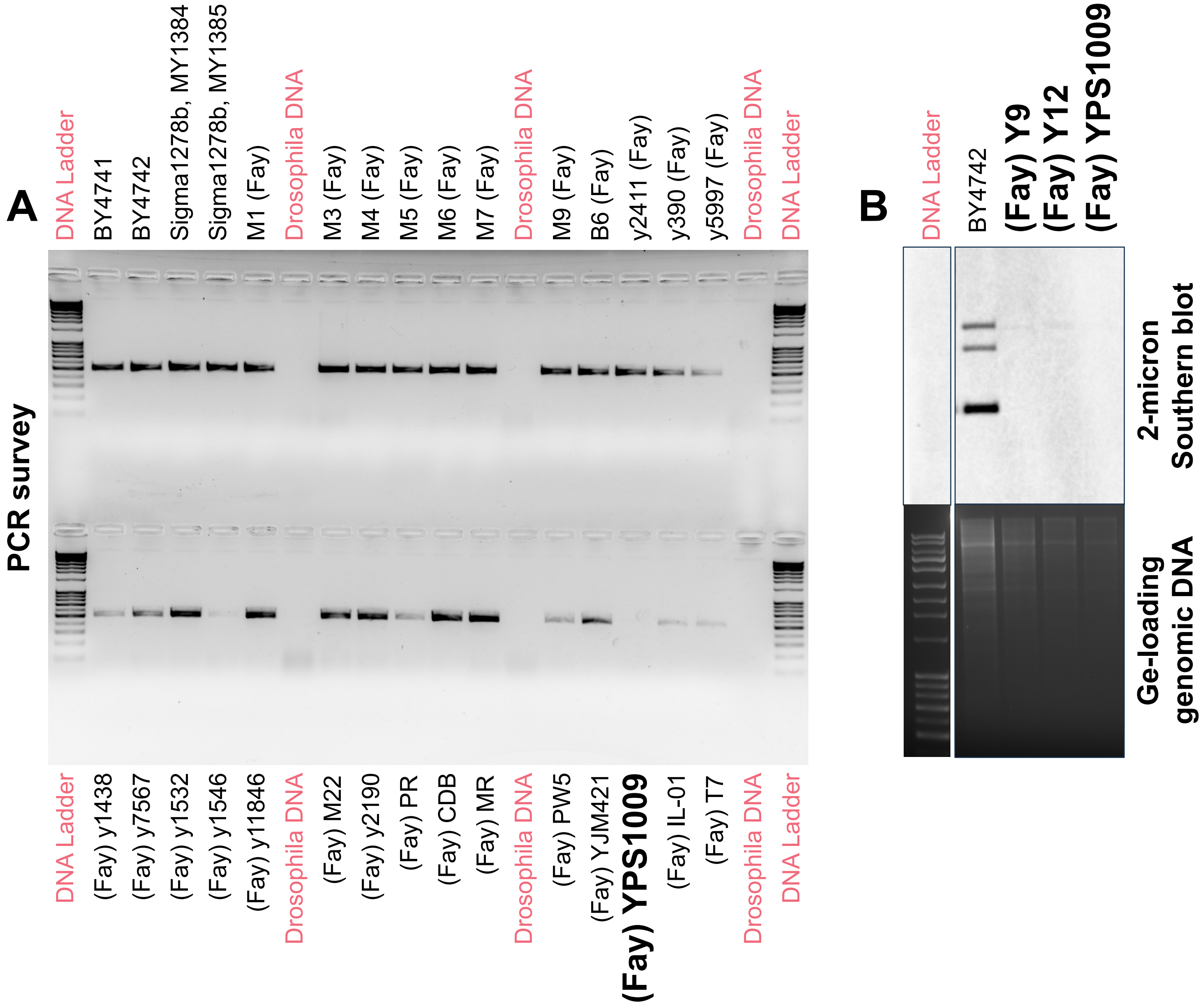


**Supplementary Figure S2**

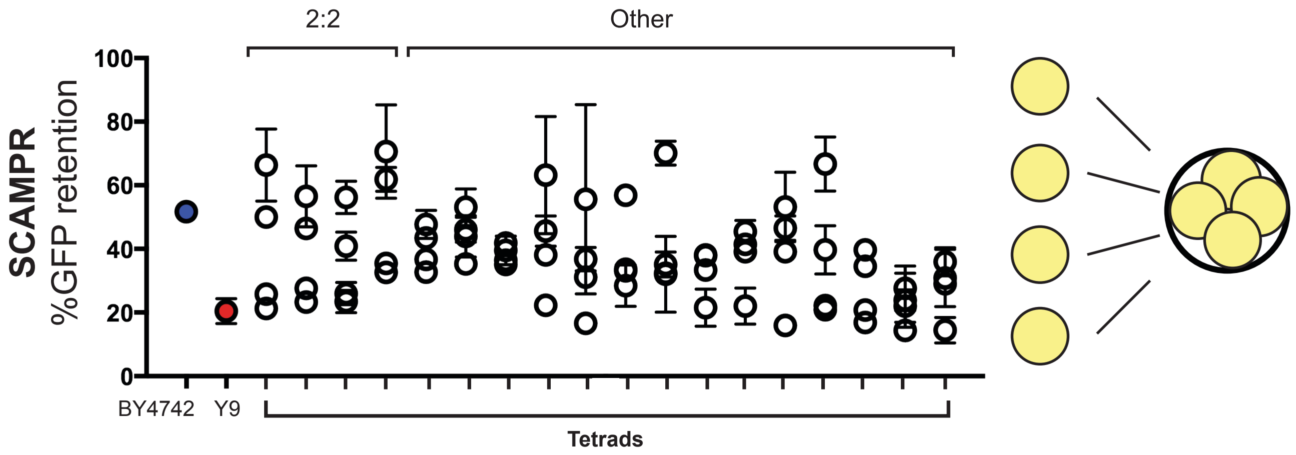


#### Supplementary Figure S3

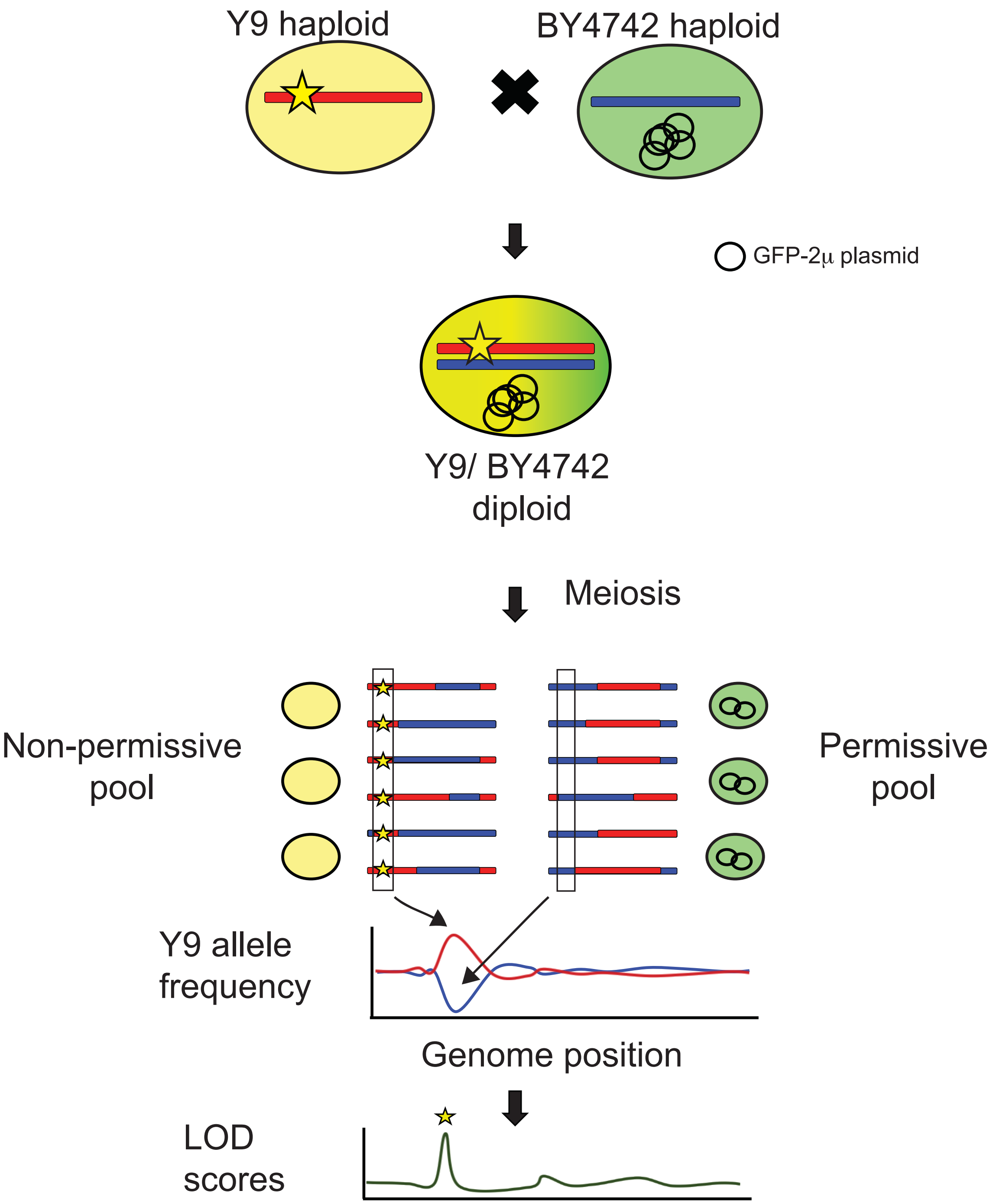


**Supplementary Figure S4**

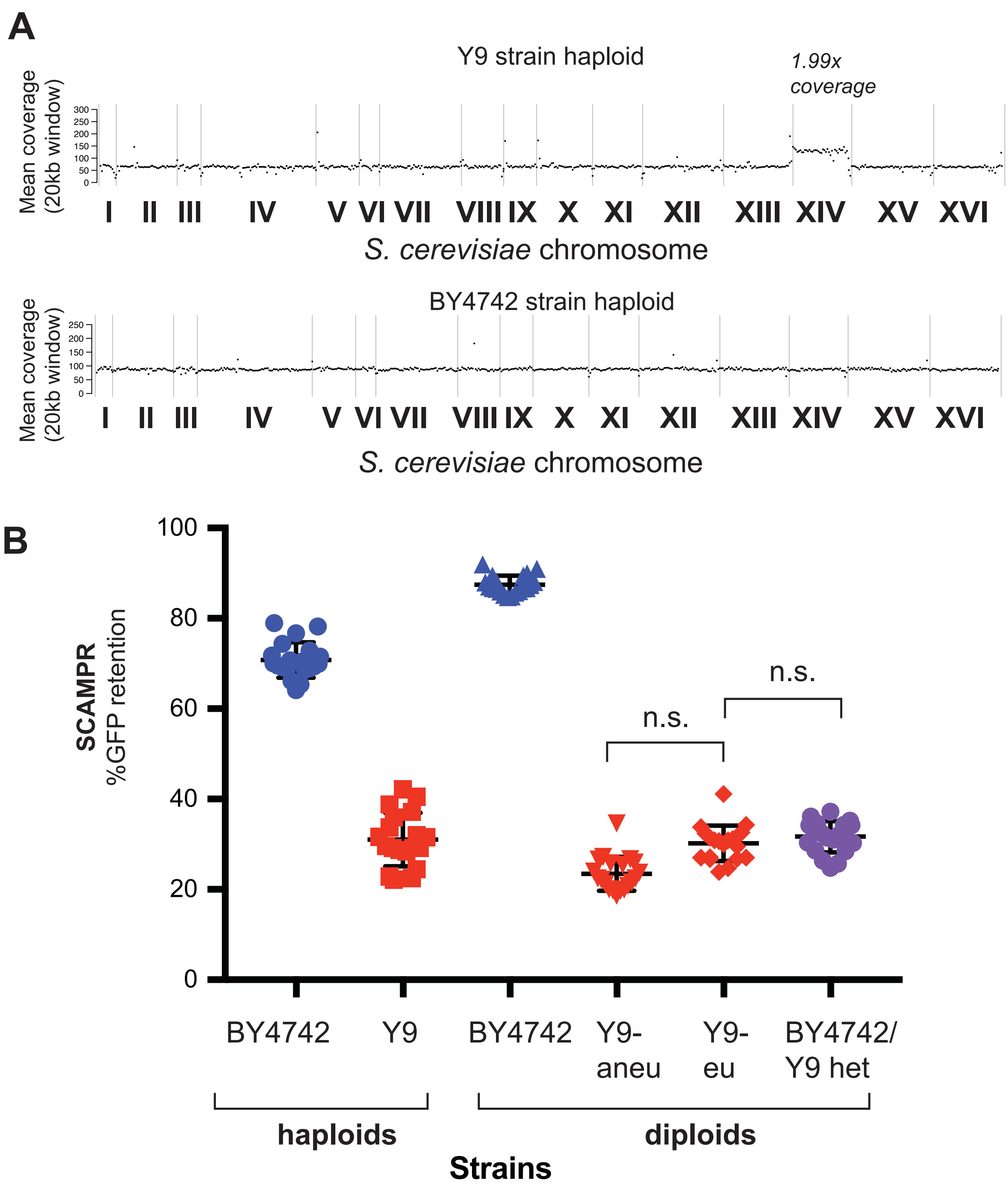


### Supplementary Figure S5

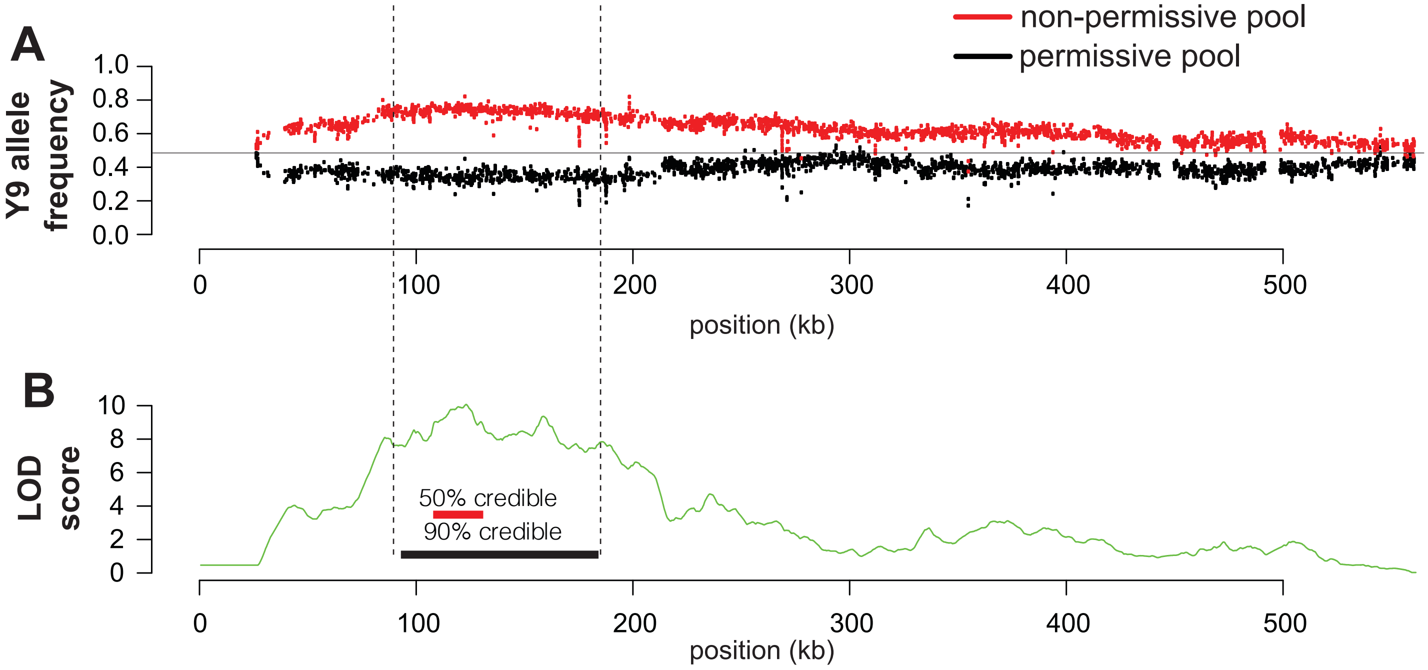


*S. cerevisiae* chromosome V

**Supplementary Figure S6**

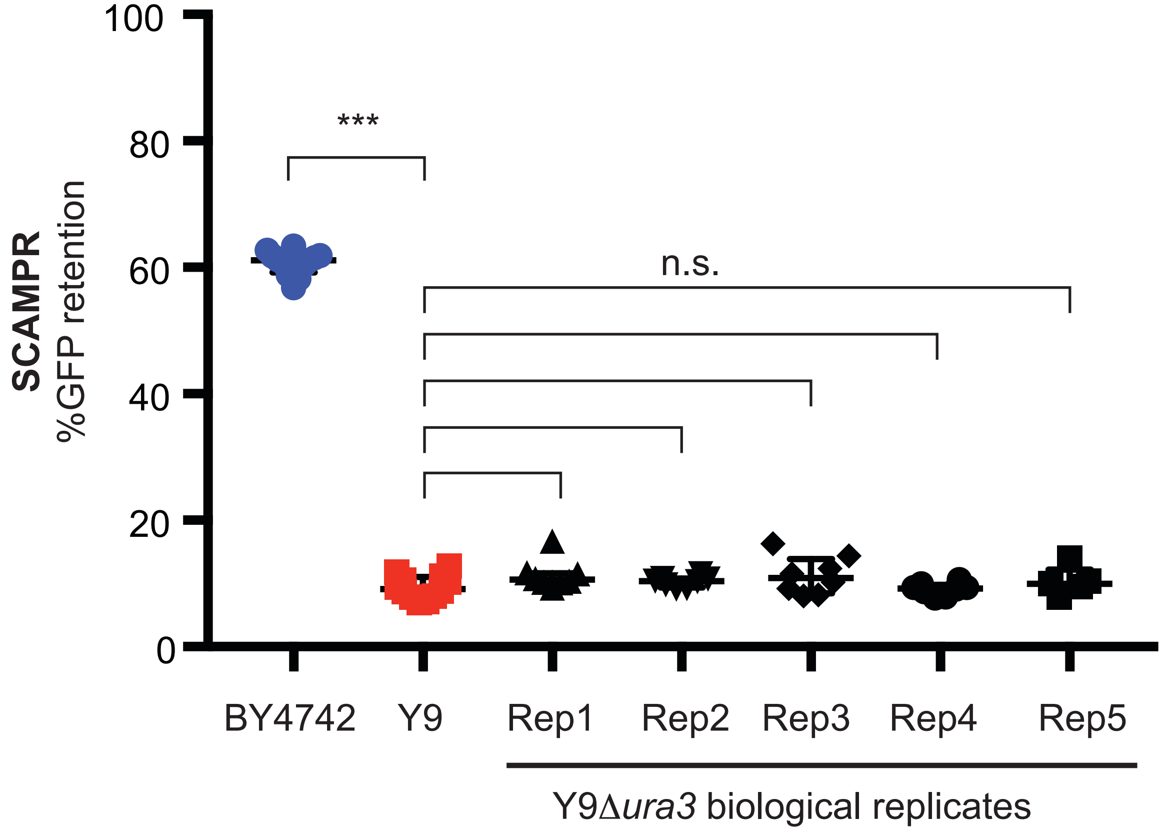


**Supplementary Figure S7**

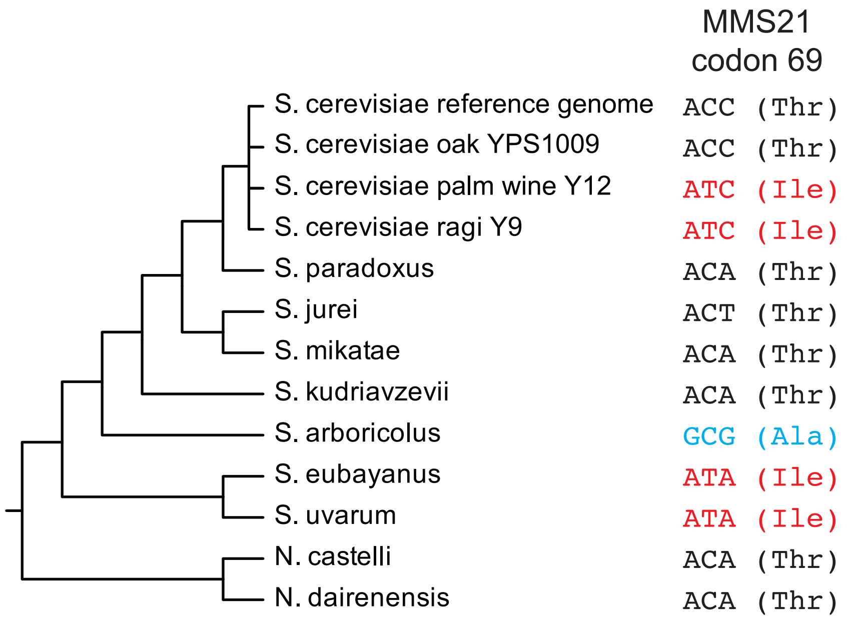


### Supplementary Figure S8

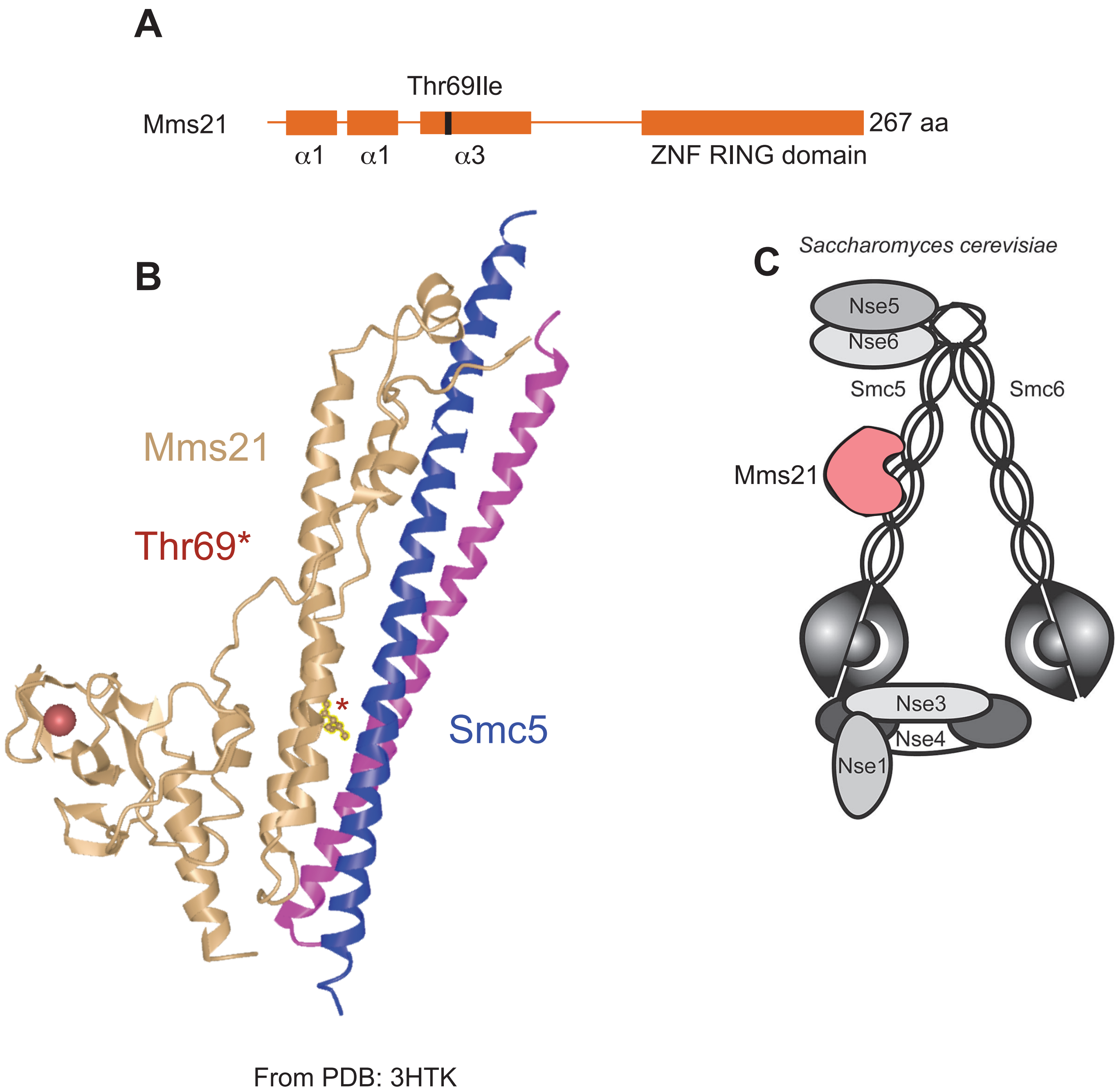


**Supplementary Figure S9**
